# A single-cell multiomics roadmap of zebrafish spermatogenesis reveals regulatory principles of male germline formation

**DOI:** 10.1101/2025.03.12.642371

**Authors:** Ana María Burgos Ruíz, Fan-Suo Geng, Gala Pujol, Estefanía Sanabria, Thirsa Brethouwer, María Almuedo-Castillo, Aurora Ruiz-Herrera, Juan J. Tena, Ozren Bogdanovic

## Abstract

Spermatogenesis is the biological process by which male sperm cells (spermatozoa) are produced in the testes. Beyond facilitating the transmission of genetic information, spermatogenesis also provides a potential framework for inter- and transgenerational inheritance of gene-regulatory states. While extensively studied in mammals, our understanding of spermatogenesis in anamniotes remains limited. Here we present a comprehensive single-cell multiomics resource, combining single-cell RNA sequencing (scRNA-seq) and single-cell chromatin accessibility (scATAC-seq) profiling, with base-resolution DNA methylome (WGBS) analysis of sorted germ cell populations from zebrafish (*Danio rerio*) testes. We identify major germ cell types involved in zebrafish spermatogenesis as well as key drivers associated with these transcriptional states. Moreover, we describe localised DNA methylation changes associated with spermatocyte populations, as well as local and global changes in chromatin accessibility leading to chromatin compaction in spermatids. Notably, we identify loci that evade global chromatin compaction, and which remain accessible, suggesting a potential mechanism for the intergenerational transmission of gene-regulatory states. Overall, this high-resolution atlas of zebrafish spermatogenesis provides a valuable resource for studying vertebrate germ cell development, evolution, and epigenetic inheritance.

## INTRODUCTION

Spermatogenesis is a biological process, which in animals occurs continuously throughout adult reproductive life. During spermatogenesis, mature haploid sperm cells (spermatozoa) are produced through a series of differentiation events from diploid spermatogonial stem cells^1^. While it was initially believed that sperm solely provided its genetic material to the egg, myriad recent studies have revealed that environmental factors, such as poor diet, exposure to toxins, and stress, can disrupt gene regulatory marks in sperm thereby influencing offspring traits^2-6^. Understanding the process of spermatogenesis and its regulatory principles is thus of crucial importance for a thorough understanding of animal reproductive processes and infertility, with applications in assisted reproductive technologies, animal breeding, and diagnostics of diverse genetic, and potentially epigenetic conditions. Single-cell sequencing technologies have enabled the generation of precise high-resolution atlases of human, mice, and primate spermatogenesis revealing both convergent and divergent pathways of spermatogenesis, each defined by their own markers, meiotic regulators, and specific ratios of germ cell populations^7-12^. Additionally, recent epigenome profiling studies have revealed that transcriptional changes during mammalian spermatogenesis are by and large paralleled by changes in chromatin accessibility^13^, 3D genome structure^14,15^ and that the initiation of meiosis is characterised by global DNA demethylation ^16^. Importantly, this global DNA methylome reprogramming event observed in spermatocytes was also observed during murine spermatogenesis ^13^ thus indicating a degree of evolutionary conservation in spermatogenesis-associated chromatin regulation. To date, the majority of genomic and imaging data of non-mammalian vertebrate (anamniote) spermatogenesis have been generated in the zebrafish (*Danio rerio*) teleost model ^17-19^. Mammalian and zebrafish spermatogenesis share core processes but exhibit differences in regulation, cellular dynamics, and organisation, the most important of which is the anatomy of the testis itself. In zebrafish, testes are organised into discrete cysts where spermatogenesis occurs synchronously within each cyst ^20^. In mammals, on the other hand, spermatogenesis occurs in seminiferous tubules with radial organisation whereas germ cells are organised in layers that progress from spermatogonia located near the basal membrane to mature sperm in the lumen ^21^. Other major differences observed in spermatogenesis between zebrafish and mammals include modes of paracrine and endocrine signalling, spermatogenesis duration, and sperm anatomy. A recent single-cell RNA sequencing (scRNA-seq) study conducted on a single biological replicate provided a useful snapshot of germ cell populations present in the zebrafish testis ^22^, whereas other high-resolution genomics work described how aging ^23^ and environmental factors ^24,25^ impact on zebrafish spermatogenesis. However, detailed genome-scale descriptions of the regulatory dynamics of anamniote spermatogenesis are still lacking, hindering our full understanding of this crucial process. To generate high-resolution transcriptional and gene-regulatory atlases of zebrafish spermatogenesis, we employed scRNA-seq and single-cell chromatin accessibility (scATAC-seq) profiling in biological replicates and combined these datasets with low input whole genome bisulfite sequencing (WGBS) of germ cell populations sorted from zebrafish testes. In our work, we provide a detailed atlas of zebrafish male germ cell types, each characterised by multiple novel transcriptional drivers. Furthermore, we identify localised DNA methylation remodelling in spermatocytes as well as transitions in chromatin accessibility leading to global chromatin compaction during spermatogenesis. Finally, we characterise in detail the chromatin makeup of elongated spermatids thus providing insight into loci with potential for intergenerational transmission of gene-regulatory states.

## RESULTS

### Cell-type resolved transcriptomes of the zebrafish testis

To generate comprehensive transcriptional and regulatory maps of zebrafish spermatogenesis, we obtained single cell suspensions from testes of adult males (*n* = 2) and prepared scRNA-seq and scATAC-seq libraries compatible with the 10X Genomics platform (**Figure 1A**). For RNA-seq, we sequenced a total of 8,432 cells from two biological replicates (**Supplementary Figure 1A** and **1B**). After initial filtering for the number of expressed genes and percentage of mitochondrial reads, both replicates displayed comparable count profiles and unsupervised cluster numbers. The majority of cells belonged to germ cell populations, however, small populations of Leydig, Sertoli, and peritubular myoid cells, were also identified (**Supplementary Figure 1B**). Given the high concordance of the two datasets, after excluding clusters exclusive to a single replicate and selecting only germ cell clusters, we integrated both replicates to obtain a total of 6,755 cells (**Figure 1B**). After manual curation of the clusters, which was based on expression of previously defined marker genes ^19,22^, we annotated major germ cell populations: undifferentiated spermatogonia-A (SPG-Aun), differentiated spermatogonia-A (SPG-Ad), spermatogonia B (SPG-B), primary spermatocytes (SPC-I), secondary spermatocytes (SPC-II), round spermatids (SPT-r), and elongated spermatids (SPT-e) (**Figure 1B and 1C**; **Supplementary Figure 1C**). We next assessed the total numbers of cells within each population and found comparable numbers of major populations between biological replicates (**Figure 1D**), as described previously^23^. Moreover, we observed that the spermatogenesis process is paralleled by a gradual transcriptional shutdown, with elongated spermatids being almost entirely transcriptionally quiescent (**Supplementary Figure 1D)**. Following annotation, we proceeded to identify cell-type-specific markers in zebrafish. This analysis, based on the newly assigned cell type identities, besides identifying previously described germline genes, identified dozens of novel cell-type specific markers of zebrafish spermatogenesis (**Figure 1E**; **Supplementary Table S1**). We next assessed gene ontology enrichments corresponding to the identified marker genes, which revealed categories in accord with their cellular function (**Figure 1F**; **Supplementary Figure 1E**; **Supplementary Table S2)**. For example, SPG populations were enriched in terms associated with ribosome biogenesis and translation, further supporting the findings that transition from self-renewal to germline differentiation is dependent on ribosome biogenesis and increased protein synthesis ^26^. SPC cells were enriched in terms associated with meiosis and DNA repair, ^21^, whereas SPTs displayed enrichment in terms linked to cilium assembly ^27^. Interestingly, the SPG-A population was enriched in categories such as: “chromatin remodelling” and “chromatin organisation”, indicative of chromatin structure changes taking place during spermatogonial differentiation.

**Figure 1).**
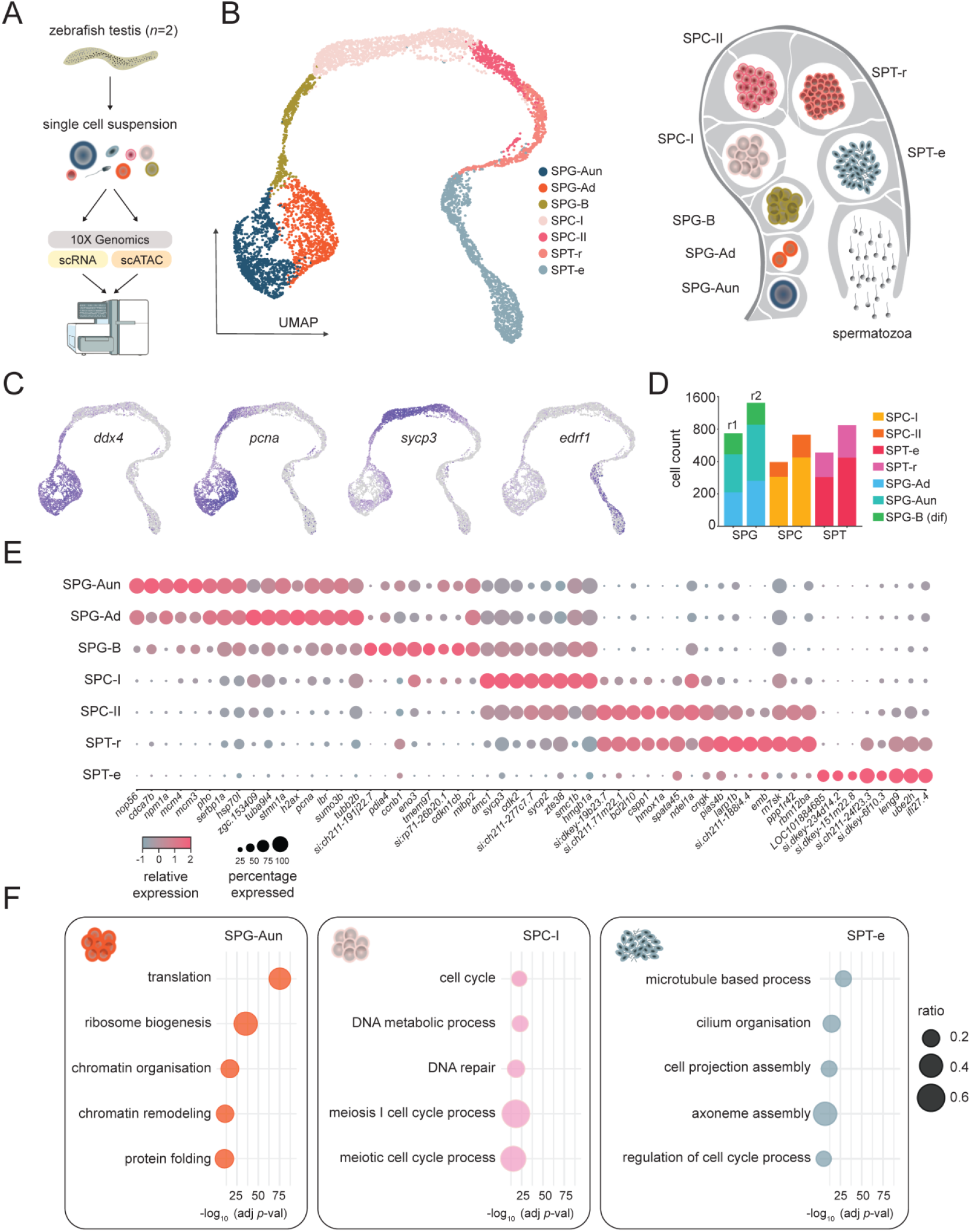
Single-cell RNA-sequencing (scRNA-seq) of the zebrafish testis. **A)** Schematic representation of experimental design and sequencing strategy. **B)** Left panel - UMAP (uniform manifold approximation and projection) plots of the zebrafish testis tissue with annotated cell types: undifferentiated spermatogonia-A (SPG-Aun), differentiated spermatogonia-A (SPG-Ad), spermatogonia B (SPG-B), primary spermatocytes (SPC-I), secondary spermatocytes (SPC-II), round spermatids (SPT-r), and elongated spermatids (SPT-e). Right panel - schematic drawing of the spermatogenesis process in *Danio rerio*. **C)** Examples of marker gene expression: *ddx4* (SPG), *pcna* (proliferation), *sycp3* (SPC), *edrf1* (SPT). **D)** Number of cells per cell type across both biological replicates (r1 – replicate 1, r2 – replicate 2). **E)** Average expression level and percentage of cells expressing each marker gene. **F)** Most highly significant gene ontology processes associated with marker genes for SPG-Aun, SPC-I, and SPT-e. Log transformed (-log10) adjusted *P* values (FDR) are represented on *x* axes.

Having identified marker genes that recapitulate germ cell-specific patterns of expression, we next wanted to search for major spermatogenesis drivers, genes that exhibit dynamic expression patterns, which reflect temporal or progressive changes during spermatogenesis. To that end, we applied an unsupervised approach for inferring linear developmental chronologies from scRNA-seq data ^28^ and identified 162 genes (**Figure 2A**; **Supplementary Figure S2A**; **Supplementary Table S3**). Moreover, to validate our driver gene set we applied an alternative approach for trajectory inference which combines dimensionality reduction with graph-based methods ^29^ (**Figure 2B**). This approach resulted in a somewhat less conservative gene population (*n* = 608), 97% of which overlapped our initial driver set (**Supplementary Figure S2B**). We next proceeded to experimentally validate drivers corresponding to major germ-cell populations by fluorescent *in situ* hybridization. We chose three novel drivers: *setb, hmgb1b*, and *ckba* that displayed distinct expression profiles corresponding to early, mid, and late spermatogenesis stages, respectively (**Figure 2C**; **Supplementary Figure S2C**). All three targets were characterised by strong staining in germ-cells with *setb* signal coinciding with *ddx4*, a canonical spermatogonial marker ^19^, and *ckba* co-staining with *sumo1*, a marker of later spermatogenesis stages^30^. Expectedly, *hmgb1b* displayed an intermediate staining profile in line with its expression pattern in spermatocytes (**Figure 2D**). Overall, our single-cell transcriptomes of the zebrafish testis provide a comprehensive overview of zebrafish spermatogenesis, delineate novel cell-type specific driver genes, and provide insight into cellular function.

**Figure 2).**
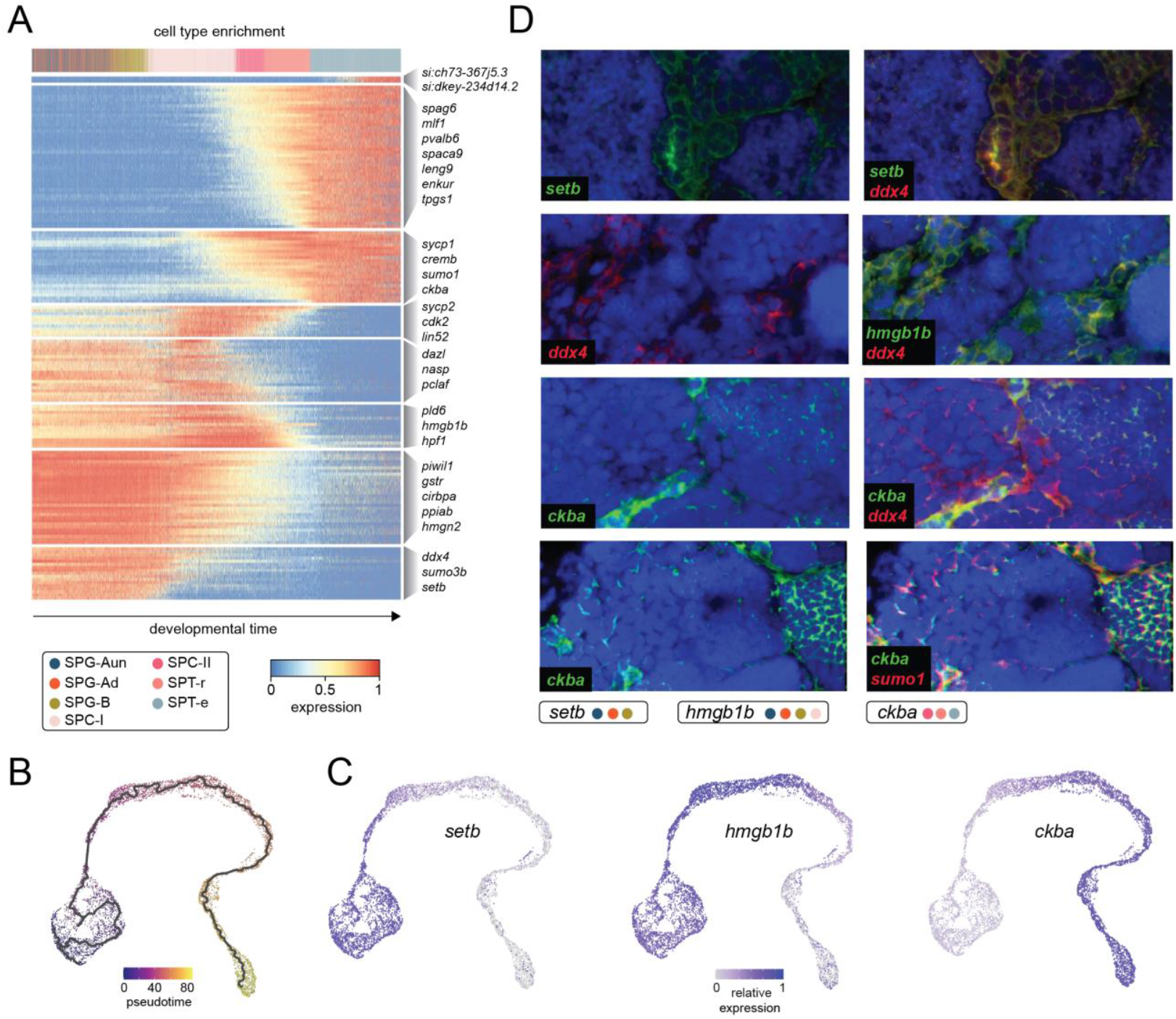
Driver genes of zebrafish spermatogenesis. **A)** Heatmap showing gene (*n* = 158) expression dynamics, with cells ranked according to their position in an inferred trajectory using SCORPIUS. Cells are color-coded by cell type, with key genes highlighted in clusters corresponding to trajectory-driven modules. **B)** UMAP (uniform manifold approximation and projection) plots of the zebrafish testis tissue denoted by pseudotime and trajectory using Monocle3. **C)** Expression pattern of newly identified spermatogenesis driver genes (*setb, hmgb1b* and *ckba*), and **D)** their double fluorescent *in situ* validation against previously defined markers (*ddx4, sumo1*).

### Localised DNA methylation changes characterise spermatocyte formation

To provide insight into gene-regulatory processes taking place during spermatogenesis, we next studied DNA methylation (5mCG), a major gene-regulatory mark required for spermatogenesis^31,32^. Moreover, 5mCG is stably maintained throughout the anamniote life cycle thereby offering a potential template for paternal epigenetic inheritance^33-36^. To that end, we generated base resolution DNA methylome (WGBS) datasets from germ cell populations sorted from zebrafish testes (**Figure 3A**). We obtained SPG, SPC-I, SPT-r populations, as well as mature sperm (SP) (**Figure 3B** and **C**). To investigate whether zebrafish spermatogenesis is characterized by global DNA methylome reprogramming processes akin to those observed in eutherian mammals^13,16^, we first assessed global 5mCG levels in these cell populations and found that the DNA methylome is stably maintained throughout spermatogenesis (**Figure 3D**). Similarly, we observed strong correlation (*r* = 0.94 - 0.96), between distinct spermatogenesis stages when 5mCG levels were compared in 10 kb genomic blocks (**Figure 3E**; **Supplementary Figure 3A**). To identify genomic regions exhibiting localised changes in 5mCG, we analysed DNA methylome profiles to detect differentially methylated regions (DMRs) > 100 bp long and with a minimum change in the fraction of methylated CpG sites (ΔmCG) of 0.2 (*P* < 0.05, Wald test). This approach revealed 3,879 localized changes in DNA methylation occurring during zebrafish spermatogenesis. *K*-means clustering (*k* = 4) revealed different clusters all of which were predominantly characterised by changes in 5mCG states associated with the SPC-I population (**Figure 3F** and **3G**). Those involved both hypo- and hypermethylated regions in SPC-I. To obtain further insight into the genomic context of these changes, we annotated the DMRs according to their genomic location ^37^ (**Supplementary Table S4, Supplementary Figure 3B**) and identified a highly significant enrichment in CpG islands (CGIs) (**Figure 3H**; **Supplementary Table S5**). CGIs are genomic regions with elevated CpG density and GC content that frequently coincide with vertebrate gene promoters and other gene-regulatory elements ^38,39^. In agreement with these results, the identified DMRs displayed strong bioCAP (CFP1-CxxC domain enrichment) signal ^40^ indicative of their CGI colocalization and their regulatory function (**Figure 3I)**. Overall, our DNA methylome data sets reveal thousands of localised, bi-directional 5mCG changes at potential gene regulatory regions occurring during spermatocyte stages, a time window, which in eutherian mammals is characterised by large-scale 5mCG remodelling ^13^.

**Figure 3).**
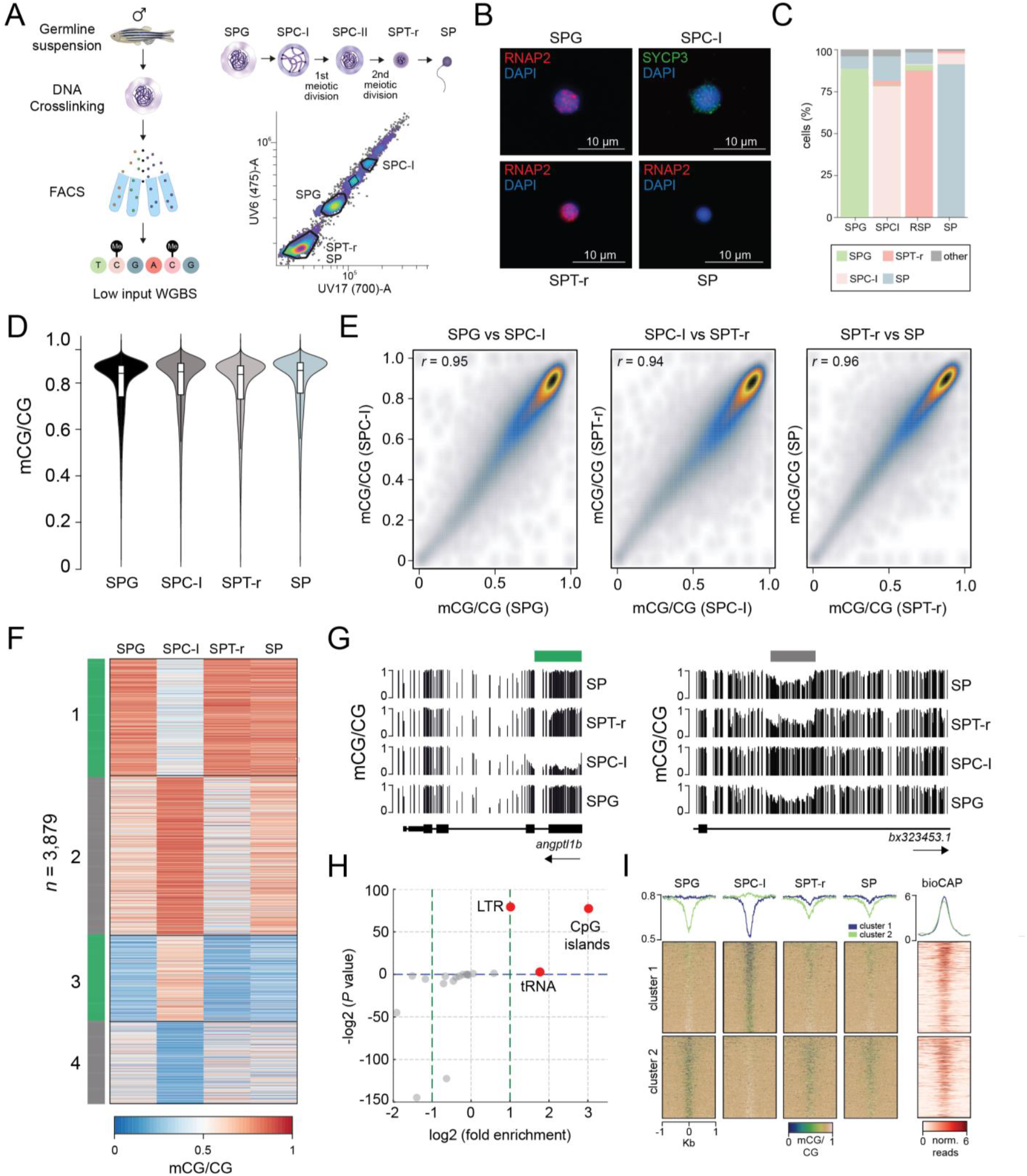
Base resolution DNA methylomes (WGBS) of zebrafish spermatogenesis. **A)** Schematic representation of the germ extraction and sequencing protocol and the sorted cell populations (SPG – spermatogonia, SPC-I – spermatocytes I, SPT-r – round spermatids, SP – mature sperm). **B)** Representative immunofluorescence images displaying DAPI-stained DNA along with specific proteins for diverse cell populations. DAPI (blue), SYCP3 (green), RNAP2 (red). **C)** Percentage of different cell populations per isolated fraction. **D)** Median DNA methylation levels (mCG) in sorted cell populations. The outer shape represents the kernel density estimate, while the boxplot inside highlights the median, interquartile range (IQR), and whiskers. Whiskers extend to the smallest and largest values within 1.5 times the IQR, with points beyond this range considered outliers. **E)** Scatter plots of average 5mCG levels in 10 kb genomic bins (*r* = Pearson correlation coefficient). **F)** K-means (*k* =4) clustering of DMRs identified between SPG and SPC-I, SPC-I and SPT-r, and SPT-r and SP methylomes. **G)** representative examples of SPC-I hypo- (left panel), and SPC-I hyper-mCG (right panel) DMRs. **H)** Enrichment of genomic features associated with DMRs. Points marked in red were deemed statistically significant (*P* < 0.05, hypergeometric test). **I)** Positional heatmaps of 5mCG and bioCAP (CpG island enrichment) signal plotted over DMRs.

### Global and local dynamics of chromatin accessibility during spermatogenesis

To study gene-regulatory mechanisms that operate during spermatogenesis, we generated 10X Chromium-compatible scATAC-seq libraries from scRNA-seq matched testes tissues (**Figure 1A**). For scATAC-seq, we sequenced a total of 17,392 single cells from two biological replicates, and after filtering for doublets and low-quality cells, we obtained 2,099 cells for replicate one and 3,251 cells for replicate two. We processed our scATAC-seq data by normalizing it for sequencing depth before applying dimensionality reduction. Using UMAP, we visualized the relationships between cells, revealing distinct populations, which were annotated by computing differential gene activity scores, predicted from the number of ATAC-seq fragments between clusters (**Figure 4A** and **4B**; **Supplementary Table 6**). After assessing the concordance of biological replicates (**Supplementary Figure 4A**), we merged the data in order to obtain a single collection of 5,350 cells, which was used for downstream analysis. The annotation was further refined by identifying marker genes linked to essential biological processes in spermatogenesis (**Supplementary Figure 4B** and **4C**). For instance, undifferentiated spermatogonia (SPG-un) were classified based on the presence of spermatogonial stem cell markers such as *id4, lin28*, and *nanos1* ^41-44^. In contrast, differentiated spermatogonia (SPG-d) were identified by the absence of these markers and the activity of genes, including *smad6, socs2, nop56*, and *zranb2*, which were previously associated with later stages of spermatogonia and their progression ^7,9,45,46^. Spermatocytes I (SPC-I) were characterized by high-level activity of genes such as *sycp3, esco2, majin* and others, which are factors essential for the early stages of meiosis, where their main role is the facilitation of homologous chromosome pairing, synapsis, and sister chromatid cohesion ^47-49^. In contrast, spermatocytes II (SPC-II) displayed chromatin opening of canonical spermatocyte markers (*spo11, mei4, tdrd12*) ^50-52^, mitotic checkpoint genes (*bub3, bub1ba*) ^53^, and genes involved in sperm motility (*foxj1* and *cfap20*) ^54,55^. Following meiosis, round spermatids (SPT-r) were distinguished by activity of *dnah1, tcte1*, and *izumo1*, which collectively guide the initial steps of spermiogenesis ^56-58^. Finally, elongated spermatids (SPT-e) were marked by the upregulation of genes such as *theg, iqcg*, and *tssk6*, central to the terminal remodeling events in spermatogenesis ^59-61^.

**Figure 4).**
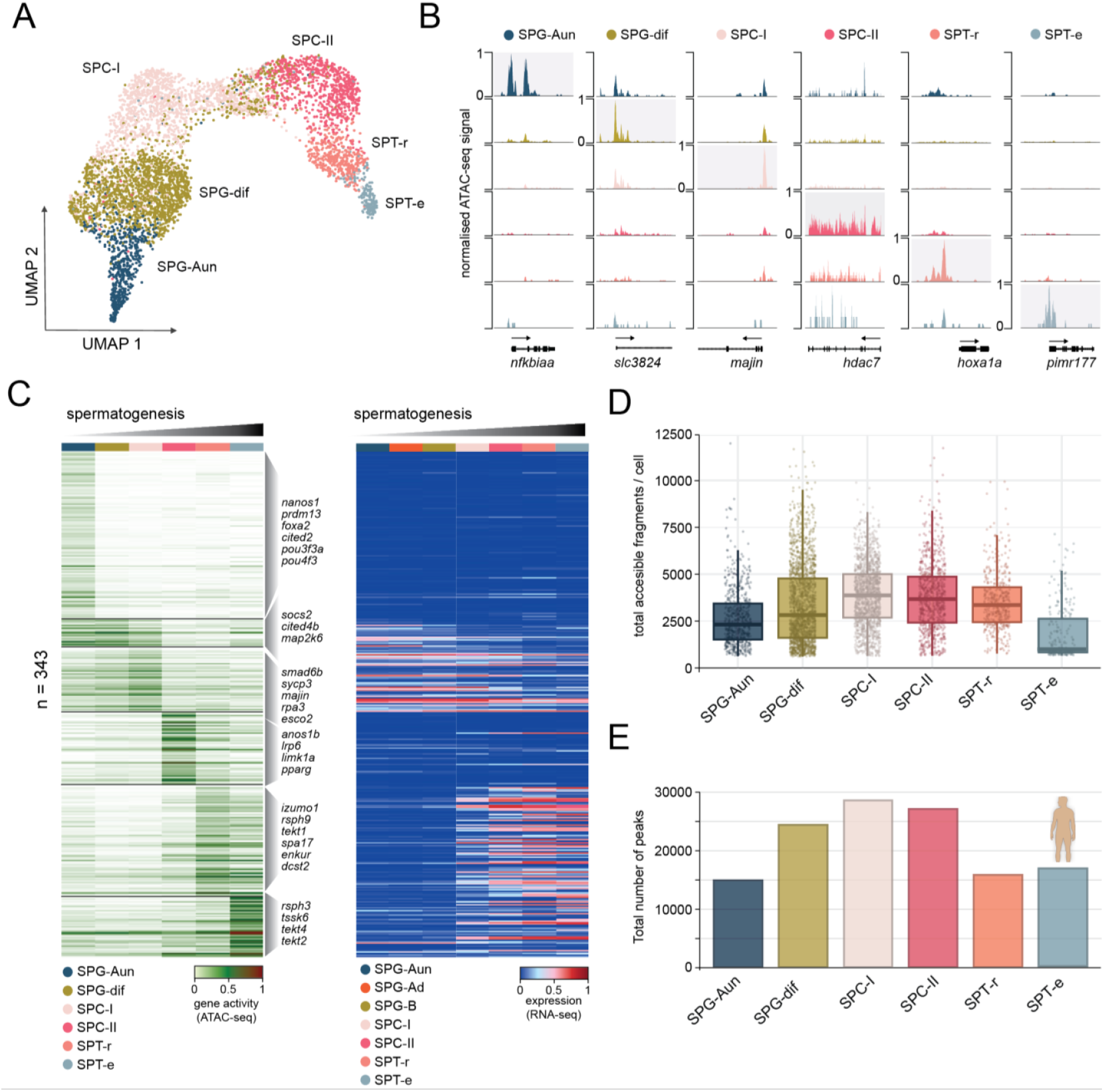
Single-cell chromatin accessibility dynamics during zebrafish spermatogenesis. **A)** UMAP (uniform manifold approximation and projection) plots of scATAC-seq data obtained zebrafish testis tissue. The annotated cell types are: undifferentiated spermatogonia-A (SPG-Aun), differentiated spermatogonia A and B (SPG), primary spermatocytes (SPC-I), secondary spermatocytes (SPC-II), round spermatids (SPT-r), and elongated spermatids (SPT-e). **B)** Genomic examples of stage- and locus-specific chromatin accessibility across identified cell populations. **C)** Heatmaps of gene activity calculated from scATAC-seq data (left panel) and matching scRNA-seq data (right panel), for genes identified as differentially active based on scATAC-seq data (log2FC > 0.8, adjusted p < 0.005), across clusters. **D)** Boxplots showing the distribution of total accessible fragments across different cell types. **E)** Total number of ATAC-seq peaks from comparable human populations identified through scATAC-seq profiling ^62^.

To understand the extent to which chromatin accessibility changes are paralleled by changes in transcription during spermatogenesis, we selected a subset of markers (*n* = 343) that displayed highly significant changes in chromatin accessibility (log2FC > 0.8 and *P* val adj < 0,005) (**Figure 4C**) and queried their transcriptional profiles. We observed a clear shift in transcriptional states coinciding with the SPC-I population that was characterised by the shutdown of spermatogonial markers and an increased expression of genes implicated in spermatocyte and spermatid formation. Having completed cluster annotation, we next studied global patterns of chromatin accessibility during spermatogenesis to better understand how global chromatin structure correlated with observed transcriptional and epigenetic changes. We found that the number of total accessible fragments per cell increased progressively from undifferentiated spermatogonial stem cells to differentiated spermatogonia (**Figure 4D**; **Supplementary Figure 4D**). This trend continued through meiosis, with peak numbers rising in spermatocytes I and II, indicating robust transcriptional activity during these stages. However, the total numbers of accessible fragments declined in round and elongated spermatids, consistent with global transcriptional shutdown and chromatin condensation. These findings are consistent with the global pattern of chromatin accessibility changes observed through single-cell approaches during human spermatogenesis.^62^ (**Figure 4E**). Overall, our single-cell chromatin accessibility datasets reveal local chromatin changes associated with activity of spermatogenesis marker genes, as well as gradual reprogramming of global chromatin structure leading to chromatin compaction and transcriptional shutdown.

### Open chromatin peaks coincide with sites of multivalent chromatin in sperm

In zebrafish sperm, “placeholder” nucleosomes and multivalent chromatin states mark thousands of regulatory regions, potentially maintaining them in a poised configuration for rapid activation in the embryo during zygotic genome activation (ZGA) ^63^. While it is believed that these specialized chromatin features could confer inherited epigenetic information and facilitate early developmental processes, it is not yet clear to what extent these regions remain in an open conformation, indicative of TF binding. To address this, we employed scATAC-seq to examine the chromatin accessibility profiles of elongated spermatids, which are expected to exhibit predominantly condensed chromatin. We identified 2,023 ATAC-seq peaks (**Supplementary Table S7**) in the SPT-e population, suggestive of some maintenance of open chromatin states during later stages of spermatogenesis. Genes associated with these peaks were enriched for several diverse biological processes, including “chromatin regulation”, “cell cycle”, and “embryonic development” (**Supplementary Figure 5A**). We next analysed the transcription factor (TF) binding motifs enriched at sites of open chromatin in elongated spermatids and found a strong association with CGI-bound (NF-Y) ^64^ and methylation-sensitive (SP1, ZBTB14, NRF1, YY2) ^38,65,66^ TFs (**Figure 5A**). Notably, this pattern was conserved throughout all spermatogenesis stages (**Supplementary Figure 5B, Supplementary Table S8**), with most enriched motifs containing a CpG site, which are otherwise strongly depleted in vertebrate genomes ^67^. Importantly, many of these factors, as well as other key components of CGI chromatin, were strongly expressed in mature sperm and zebrafish testis tissues, based on RNA-seq data re-analysed from previous studies ^34,68^ (**Figure 5B**). To clarify the relationship between ATAC-seq signal and DNA hypomethylation, a common feature of CGIs, we *de novo* identified hypomethylated CpG-rich regions (unmethylated regions – UMRs) ^69^ using our germ cell stage-specific DNA methylation data (**Figure 3**). These analyses revealed similar numbers of UMRs throughout spermatogenesis (spermatogonia = 18,406; mature sperm = 18,058) (**Figure 5C**), in line with the absence of any major DNA methylome reprogramming events (**Figure 3**). Next, we plotted ATAC-seq signal from our single-cell data across mature sperm UMRs, revealing a chromatin reprogramming pattern analogous to the one observed on the genome-wide scale (**Figure 5D**; **Supplementary Figure 5C**). Notably, peak intensity and width both varied, with SPG-dif and SPC-I peaks being the broadest. Given the strong correlation between ATAC-seq signal and DNA hypomethylation (**Figure 5D** and **5E**), we next asked whether these sites overlap with multivalent and “placeholder” chromatin regions (enriched in H3K4me1, H3K4me3, H2AZ, H3K14ac, and hypomethylation) (**Figure 5F**) ^63^. Indeed, late SPT-e ATAC-seq signal strongly coincided with placeholder chromatin, previously identified as a major driver of maternal-to-paternal chromatin remodeling before ZGA. Taken together, our findings suggest that thousands of open chromatin regions might be retained intergenerationally within the context of placeholder chromatin, likely to facilitate ZGA and early embryonic development.

**Figure 5).**
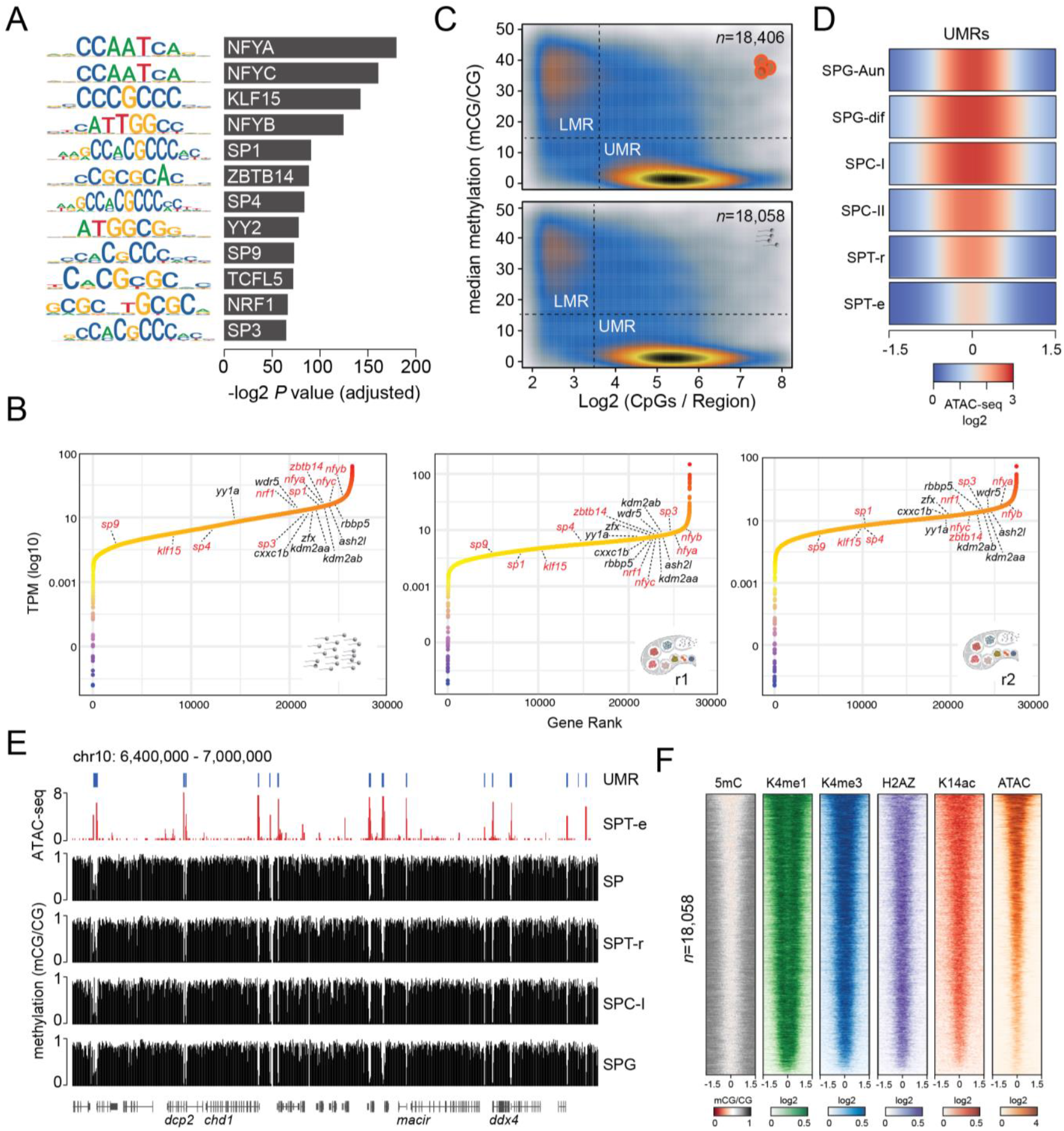
Chromatin state of elongated spermatids (SPT-e). **A)** Top enriched TF binding motifs in open chromatin peaks retained in SPT-e, ranked in descending order (*P* value < 0.05; hypergeometric test. **B)** Gene expression (transcripts per million, TPM) plotted against gene rank, in transcriptomes of mature zebrafish sperm (left panel) ^31^ and two biological replicates of zebrafish testis (middle and right panel) ^64^. Transcripts coding for TFs that display enriched motifs in SPT-e are marked in red. Other key components of CGI chromatin are highlighted in black. **C)** *De novo* discovery of regulatory regions (UMRs – unmethylated regions - i.e CGI promoters; and LMRs – lowly methylated regions – i.e enhancers). Scatter plot representing CpG density plotted against DNA methylation levels in newly identified UMRs and LMRs. Upper panel: spermatogonial UMRs (*n* = 18,406), lower panel: mature sperm UMRs (*n* = 18,058). **D)** scATAC-seq signal (log2) of germ-cell populations plotted over mature sperm UMRs (*n* = 18,058). **E)** Genomic example of a 600 kb region surrounding the *ddx4* gene demonstrating the concordance between DNA methylation, UMRs and scATAC-seq signal (SPT-e). **F)** Positional heatmaps of i) mature sperm DNA methylation (this study), ii) H3K4me1, H3K4me3, H2AZ, and H3K14ac, ChIP-seq enrichment signal from mature sperm ^63^, and iii) scATAC-seq signal from SPT-e, plotted over UMRs.

## DISCUSSION

The process of spermatogenesis is largely conserved among vertebrates and is generally divided into three phases: spermatogonial proliferation, meiosis, and post-meiotic maturation. Numerous studies, including recent single-cell genomics reports, have greatly advanced our understanding of the cellular population dynamics and gene-regulatory events that occur during spermatogenesis ^7,8,11-13,70^. However, most of this research was conducted on mammalian models, leaving a significant gap in our knowledge of the regulatory mechanisms governing non-mammalian (anamniote) sperm formation. This is important to appreciate, because mammals and anamniotes exhibit major differences in chromatin regulation during germline and early development. For example, mammalian sperm formation involves near-complete replacement of histones with protamines, resulting in highly condensed sperm chromatin, with only a small fraction of histones (1 - 10%) retained. Nevertheless, the extent of this phenomenon as well as the exact genomic locations of retained loci and their function, remain a topic of debate ^71-75^. In contrast, fish display diverse sperm chromatin packaging strategies. Zebrafish, for instance, predominantly use histones to package their sperm ^76^, whereas medaka employ protamine-based packaging ^77^, similar to mammals The presence of nucleosomes in vertebrate sperm suggests a potential mechanism for intergenerational epigenetic marking, which may be even more pronounced in species like zebrafish that rely on nucleosome-based sperm packaging. To investigate the gene-regulatory processes involved in zebrafish sperm formation, and to better understand the potential for paternal inheritance of epigenetic states, we generated scRNA-seq and scATAC-seq datasets of the zebrafish testis. Our scRNA-seq data revealed seven major germ-cell populations corresponding to spermatogonia (SPG-Aun, SPG-Ad, SPG-B), spermatocytes (SPC-I, SPC-II), and spermatids (SPT-r, SPT-e) (**Figure 1**), as well as Leydig and Sertoli cells (**Supplementary Figure 1**). Overall, our findings are consistent with previously published zebrafish scRNA-seq datasets, though we observed minor discrepancies in cell population ratios ^22^, likely due to differences in sequencing depth or zebrafish age ^23,78^. Consistent with previous work, we also found that the number of expressed genes decreases progressively as spermatogenesis advances ^78^, resulting in global transcriptional downregulation in elongated spermatids (**Figure 6**). Finally, by inferring developmental trajectories, we identified novel driver genes across all stages (**Figure 2**), generating a resource that will be of great value for understanding infertility, and other disease states linked to spermatogenesis. Our open chromatin profiling via scATAC-seq broadly recapitulated the spermatogenesis stages identified through expression profiling, while providing a higher-resolution view of gene-regulatory changes (**Figure 4**). Differential analysis of scATAC-seq data revealed thousands of changes in chromatin accessibility, many of which paralleled changes in steady-state RNA levels. We further observed a gradual increase in chromatin accessibility during spermatogonial differentiation, peaking at SPC-I and SPC-II stages, followed by a progressive decrease in round and elongated spermatids. This pattern is consistent with Gene Ontology (GO) enrichments of marker genes expressed in differentiating spermatogonia, which are strongly associated with functions related to chromatin remodeling and organization (**Figure 1**). A similar global pattern of chromatin reprogramming was also reported during human spermatogenesis in a recent scATAC-seq study ^62^, whereas scCOOL-seq profiling of human sperm at single-cell resolution revealed a somewhat different pattern yet still exhibited open chromatin peaks during spermatocyte stages ^13^. Collectively, these observations underscore the deeply conserved nature of global chromatin state changes across vertebrates and their importance in shaping male germ cell development. To better understand the epigenomic changes that occur during zebrafish spermatogenesis, we complemented our findings with DNA methylome data obtained by WGBS from four germ cell populations. In mammals, two rounds of genome-wide DNA methylation reprogramming occur; one during pre-implantation development and another during primordial germ cell formation ^79-86^. More recently, a major DNA methylation reprogramming event specifically at the spermatocyte stage has also been reported in humans ^13,16^. However no large scale epigenome rearrangements have been observed in zebrafish to date ^35,36,87,88^ suggesting that parental inheritance of gene-regulatory marks and other epigenetic factors from gametes in anamniotes may play a more significant developmental role. Consistent with previous zebrafish DNA methylome profiling studies, we did not detect any major DNA methylation reprogramming events during spermatogenesis (**Figure 3**). Interestingly, the majority of statistically significant DNA methylation changes (*n* = 3,879; ∼0.13% genome) (**Figure 6**) that we identified, were associated either with hyper- or hypo-methylation in spermatocytes, which is when DNA methylation reprogramming takes place during spermatogenesis in mammals. Whether these changes in both mammals and anamniotes reflect genuine regulatory events or are simply byproducts of large-scale chromosomal rearrangements ^89,90^, remains an open question. Finally, using our DNA methylome data, we *de novo* identified CpG-rich unmethylated regions (UMRs; ∼3% genome) and demonstrated that these sites retain open chromatin and multivalent “placeholder” chromatin (**Figure 6**). These findings provide further evidence for intergenerational inheritance of epigenetic marks in zebrafish ^63^, suggesting that open chromatin and transcription factor binding likely play a role in this process. Several CGI - associated chromatin components (e.g., NRF1, SP5) have already been implicated in infertility and spermatogenesis defects ^91,92^, highlighting the potential relevance of CGI chromatin for intergenerational epigenetic transmission ^93^, though the full extent of this phenomenon remains to be determined. Our study thus offers an important roadmap for tackling fundamental questions in spermatogenesis regulation. For example, the provided data can inform research into stem cell maintenance and niche regulation, as well as help elucidate how chromosomal rearrangements, homolog pairing, and crossover formation influence the flow of epigenetic information. Overall, we present a significant community resource that advances understanding of vertebrate spermatogenesis, epigenetic inheritance, and the evolutionary conservation of gene regulatory processes associated with germ cell development.

**Figure 6).**
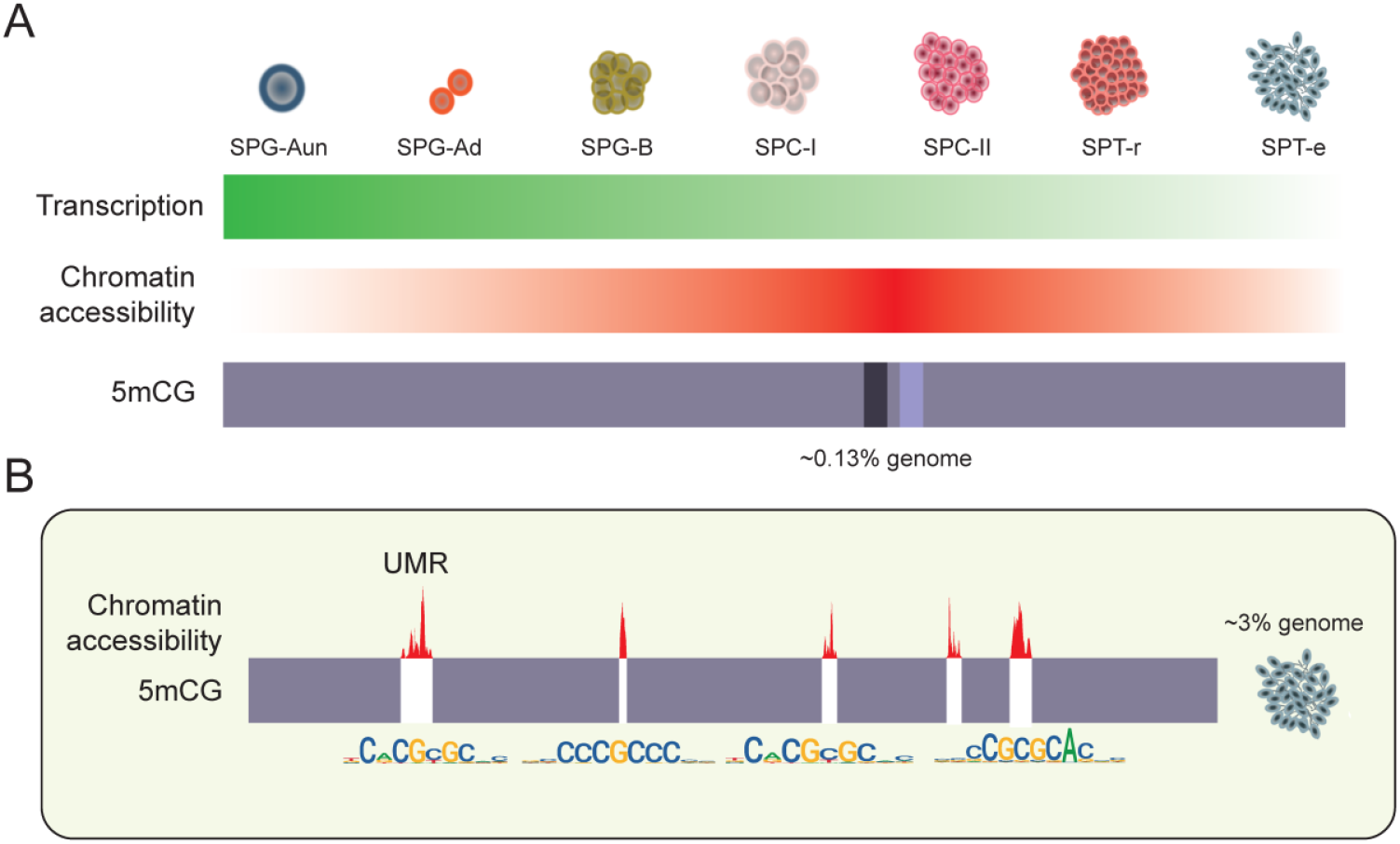
Chromatin and transcriptome dynamics during zebrafish spermatogenesis. **A)** Schematic representation of transcriptome (green), chromatin accessibility (red), and DNA methylation (purple) dynamics during zebrafish spermatogenesis. The spermatogenesis transcriptome is characterized by a gradual transcriptional downregulation, leading to the lowest transcriptional activity in elongated spermatids. Chromatin accessibility, on the other hand, gradually increases, reaching peak activity during spermatocyte development (SPC-I and SPC-II). Meanwhile, the DNA methylome remains stable, except for localized methylation and demethylation events (dark and light shading) affecting 0.13% of the genome during spermatocyte formation. **B)** Unmethylated regions (UMRs) exhibit strong ATAC-seq signal and an enrichment in CpG island (CGI) protein-binding motifs in elongated spermatids (SPT-e). This chromatin configuration affects approximately 3% of the genome, and at least a fraction of this epigenetic makeup may be inherited intergenerationally through sperm.

## MATERIALS AND METHODS

### Data availability

All 10X Genomics data (scRNA-seq and scATAC-seq), have been deposited in Gene Expression Omnibus (GEO) database, www.ncbi.nlm.nih.gov/geo (GSE283803 and GSE283804). WGBS data have been deposited in Array Express under the accession number E-MTAB-14873.

### Ethics declarations

Zebrafish experiments were approved by the Garvan Institute of Medical Research Animal Ethics Committee under AEC approval 17/22. All procedures complied with the Australian Code of Practice for the Care and Use of Animals for Scientific Purposes. Animal experiments conducted at CABD have been approved by the Animal Experimentation Ethics Committees at the Pablo de Olavide University and CSIC (license number 02/04/2018/041). Procedures at UAB complied with the animal ethics guidelines approved by University Animal Experimentation Ethics Committee.

### Zebrafish procedures

Adult transgenic zebrafish (*Danio rerio*), aged 6 months, were prepared and anesthetized with 0.25% tricaine methanesulfonate (MS-222) on ice for 15 minutes before experimentation. Testes were dissected and rinsed 3 times with PBS. The samples were then digested in 10 ml of 0.25% trypsin at 37 °C for 15 minutes, with gentle pipetting every 3 minutes to facilitate tissue breakdown. Digestion was halted by adding DMEM supplemented with 10% FBS, and the resulting cell suspension was passed through a 70 μm nylon mesh filter. The cells were then centrifuged at 1,000 rpm for 10 minutes, after which the pellet was resuspended in 1 ml of DMEM (with 10% FBS) and filtered through a 40 μm nylon mesh.

### scRNA-seq and scATAC-seq library construction and sequencing

Expression (scRNA-seq) libraries were prepared using the 10X Chromium 3′ GEM v3.0 kit and sequenced on an Illumina NovaSeq 6000 platform (S4, 200 bp, PE). Chromatin accessibility (scATAC-seq) libraries were generated using the 10X Genomics v1.0 capture kit and sequenced on an Illumina NextSeq 550 platform with the High Output Kit v2.5 (150 bp, PE). For both scRNA-seq and scATAC-seq, we obtained an average of 25,000–30,000 paired-end reads per cell.

### Single-cell RNA-Seq read alignment and quantification

Raw reads were demultiplexed and mapped to the zebrafish danRer11 reference transcriptome using the 10X Genomics CellRanger (v7.2.0) pipeline ^94^. Before downstream QC, CellRanger gene expression algorithm identified 3,984 cells in replicate 1 and 5,996 cells in replicate 2. Following filtering, these counts were reduced to 2,852 cells (replicate 1) and 5,996 cells (replicate 2), with median gene counts of 2,436 and 2,388, respectively.

### Pre-processing, quality filtering, batch integration and dimensional reduction scRNA-seq

Seurat v5.1.0 ^95^ was used for scRNA-seq preprocessing, quality control, and analysis. Raw count matrices from two 10X Genomics replicates were imported. Cells expressing < 200 or > 7000 genes or with > 5% mitochondrial content were excluded. Data were normalized using *LogNormalize* (scale factor: 10,000), and highly variable features were identified via vst. Replicates were integrated using *harmony::RunHarmony*. To identify cell populations, PCA was performed, and the optimal number of PCs was determined using ElbowPlot, selecting the first 11 PCs. A shared nearest neighbour (SNN) graph was built with *Seurat::FindNeighbors*, followed by clustering via Leiden (resolution = 0.5). Clusters with low UMI and gene counts, ribosomal RNA enrichment, or clusters present only in one replicate, were removed as likely artifacts. Additionally, clusters expressing Leydig (*cyp17a1, hsd3b1*), Sertoli (*fshr, nr5a1*), and peritubular myoid cell markers (n=219 cells) were excluded from further analyses. The analysis was repeated on the remaining cells using 6 PCs, yielding 13 clusters. Cells were visualized using UMAP (*RunUMAP*, 8 PCs).

### Cell classification and marker identification

Marker genes for each cluster were identified using *Seurat::FindAllMarkers*, applying a log fold-change threshold of 0.15 and restricting the analysis to positive markers, while excluding genes expressed in fewer than 10 cells. Clusters were manually annotated based on literature-defined marker genes ^19,22^. Spermatogonia (SPG), which comprised clusters 8, 6, 3, 5, 12, and 10, were characterized by *ddx4* and *piwil1* expression. SPG-A (clusters 8, 6, 3, 5) exhibited higher *ddx4* and *piwil1* expression compared to SPG-B (clusters 10, 12). Within SPG-A, clusters 3 and 5 (A - undifferentiated) were characterized by *eno3, e2f5*, and *ripply2*, while clusters 6 and 8 (A - differentiated) expressed *hist1h2a6*. Spermatocytes (SPC), which comprised clusters 0, 4, and 9, were identified by *sycp3* and *pcna* expression, where SPC-I (clusters 0, 4) exhibited high *sycp3* and *pcna* levels and SPC-II (cluster 9) exhibited lower expression of these markers. Spermatids (SPT) were categorized into round SPT (clusters 7, 11), which lacked *edrf1*, and elongated SPT (clusters 2, 1), which expressed *edrf1*. Upon annotation, we conducted a refined analysis by employing a higher log fold-change threshold (0.25), which led to the identification of new zebrafish cell-specific markers, thus further improving cell-type resolution.

### Gene set enrichment analysis

Gene enrichment analysis was conducted using the marker genes identified for each cell type. These genes were input into the g:Profiler website tool ^96^, with *Danio rerio* specified as the reference organism. The analysis focused exclusively on enriched biological processes associated with each cell type. The most representative terms were ranked by statistical significance, and the top five biological processes with the lowest adjusted *P* values were selected for visual representation.

### Double fluorescent *in situ* hybridization (FISH) on zebrafish testis cryostat sections

The coding sequences of *ddx4, setb, hmgb1b, ckba* and *sumo1* (**Supplementary Table S9**) were synthesized and introduced into pBluescript KS+ plasmid using *NotI* and *XbaI* restriction enzymes (GENEWIZ, Azenta Life Sciences). Antisense riboprobes were synthesized using digoxigenin-11-UTP or fluorescein-12-dUTP (Merck) and T3 RNA polymerase (Merck) after *SacI*-linearized plasmid digestion. Cryosectioning of samples was performed as previously described ^97^ followed by an adapted double FISH protocol ^98^. Sections underwent Proteinase K treatment (10 µg/mL, 30 min, 37°C) and post-fixation with 4% formaldehyde (30 min, RT). Probe hybridization occurred at 70°C for ≥ 16h. After post-hybridization SSC washes, probes were developed using anti-DIG-POD antibody (Merck, 1:150) and incubated overnight (4°C). Sections were washed (PBS, Borate buffer) before staining with green TSA amplification (50 μg/mL TSA Fluorescein) for 30 min (RT, dark). POD enzyme was quenched (0.1M Glycine, pH 2.2), followed by anti-Fluorescein-POD antibody incubation (Merck, 1:150) and overnight (4°C) incubation. After further washes, red TSA amplification (50 μg/mL Red TSA) was applied (30 min, RT, dark). Sections were washed (PBS) and incubated overnight (4°C) with DAPI (1:5000, Sigma). Confocal imaging was performed using an LSM 880 (Zeiss) and processed with ImageJ ^99^.

### scRNA-seq trajectory analysis

For trajectory analysis of scRNA-seq data, we employed Monocle3 (v1.2.9) ^28^ and SCORPIUS (v1.0.9) ^29^ R packages with imported clustering information from Seurat, including UMAP coordinates and cluster identities. The trajectory graph was constructed using the *monocle3::learn_graph* function, and cells were ordered along the pseudotime trajectory with *order_cells*. Differential expression analysis across pseudotime was performed using *monocle3::graph_test*. SCORPIUS was used to validate the findings, starting with dimensionality reduction using multidimensional scaling (MDS) in three dimensions, based on Spearman correlation distance. A trajectory was inferred with default parameters, and candidate marker genes were identified using *SCORPIUS::gene_importances* function, selecting the top 200 genes for further analysis. Genes were grouped into modules excluding ribosomal-associated modules (module < 9). To refine the results, driver genes identified by Monocle3 and SCORPIUS were filtered and compared. For Monocle3, genes were filtered by Moran’s I > 0.5 and *q*-value < 0.05, resulting in 604 driver genes. For SCORPIUS, the 162 genes identified through the *SCORPIUS::gene importances* function were extracted. A comparison between the two methods revealed an overlap of 158 genes.

### Single-cell ATAC preprocessing and quality control

Replicate data (r1, *n* = 8,305 cells; r2, *n* = 9,088 cells) were processed using the Cell Ranger ATAC (v.1.2) count pipeline (danRer11 assembly). The output of the Cell Ranger ATAC pipeline was utilised as the input for downstream analysis using the Signac R package (v1.14.0). A unified peak set was generated using the *GenomicRanges::reduce* function to merge peak coordinates from both datasets, with subsequent filtering based on peak length. Fragment objects were created for each sample using the *Signac::CreateFragmentObject* function, and peak quantification was conducted using the *Signac::FeatureMatrix* function. Cells deemed low-quality were identified and excluded if they had a nucleosome signal score below 1 and a TSS enrichment score below 5. Meanwhile, cells with more than 40% of fragments mapping to peaks and between 500 and 10,000 fragments in peaks were retained for further analysis. To identify potential doublets, scDblFinder R package (v1.8.0) was applied to the dataset using a converted SingleCellExperiment object from Seurat. Only cells classified as “singlet” were retained for downstream analyses. Following filtering, the number of retained cells was *n* = 2,099 (r1) and *n* = 3,251 (r2). After quality filtering, term frequency-inverse document frequency (TF-IDF) normalisation was applied, followed by the selection of relevant features using a threshold (min.cutoff = 0.5). Dimensionality reduction was performed using singular value decomposition (SVD) to compute latent semantic indexing (LSI) embeddings with up to 70 dimensions. To assess the impact of sequencing depth, LSI components were evaluated using *Signac::DepthCor*, and those with strong correlations to sequencing depth were excluded (LSI1 and LSI10 in replicate 1, LSI1 and LSI8 in replicate 2). The remaining components were used for downstream analyses.

### Single-cell ATAC analysis

Integration of both replicates was performed using the *Seurat::FindIntegrationAnchors* and *Seurat::Integrate-Embeddings* functions, specifying dimensions 2:6 and 9:70 for integration. Clustering was conducted using the *Seurat::FindCluster* function with the smart local moving (SLM) algorithm for modularity optimization, with a resolution parameter of 1.2 and algorithm set to 1. Clusters that were found exclusively in one of the replicates were removed. Annotation was performed by first calculating gene activity scores using the *Signac::GeneActivity* function. Annotation was further refined by identifying marker genes associated with key biological processes in spermatogenesis. Using these gene activity scores, the RNA data was log-normalised, and differentially expressed genes were identified using the MAST test, adjusting for RNA count as a latent variable. Genes with an adjusted *P* < 0.005 and avg_log2FC > 0.8 were considered significant. Motif position frequency matrices (PFMs) from the JASPAR CORE vertebrate collection were obtained using the *Signac::getMatrixSet* function and added with *Signac::AddMotfs*. Enriched motifs were identified with *Signac::FindMotifs* in peaks accessible in ≥10% of elongated spermatids.

### Flow cytometry and cell enrichment analysis

Zebrafish testes were disaggregated following a previously described protocol ^25^. The cell suspension was fixed in 1% formaldehyde, centrifuged, and stored at -80ºC until use. Frozen samples were then resuspended in PTBG (0.05% Tween-20 in 1X PBS) and immunostained in solution with a mouse anti-RNA polymerase II antibody (#ab5408, Abcam) diluted in PTBG (1:1000) and incubated at 4ºC overnight. The next day, cells were centrifuged for 15 min at 2000 g, resuspended in PTBG, and incubated for 5 min at 21ºC with an anti-mouse Cy5 (#115-175-166, Jackson ImmunoResearch, 1:1000). After a PTBG wash, cells were stained with 5 μg/ml Hoechst 33342 for 35 min at 21ºC and sorted using a BD FACS Discover S8 Cell Sorter. Four testicular populations (spermatogonia, spermatocytes I, round spermatids, and spermatozoa) were isolated considering their nucleus complexity, ploidy, and RNA polymerase II staining, with between 60,000 and 400,000 cells collected per cell type. Cell enrichment of each flow-sorted population was evaluated by immunofluorescence. Sorted cells were fixed onto slides by incubating with freshly prepared 4% paraformaldehyde solution containing 0.15% Triton X-100 for 2 hours at room temperature in a humidified chamber. After air-drying, the slides were washed in 1% Photo-Flo and incubated with the primary antibodies rabbit anti-SYCP3 (#ab15093, Abcam, 1:100) and mouse anti-RNA polymerase II (#ab5408, Abcam, 1:1000) overnight at 4ºC. After incubation, slides were washed twice in PTBG, followed by a 1-hour incubation at 37ºC with the secondary antibodies anti-rabbit FITC (#111-095-003, Jackson ImmunoResearch, 1:200) and anti-mouse Cy5 (#115-175-166, Jackson ImmunoResearch, 1:1000). DNA was counterstained with antifade solution containing 0.1 μg/ml DAPI and stored at -20ºC until use. Stained slides were analyzed using an epifluorescence microscope (Axiophot, Zeiss). Spermatogonia (SPG) exhibited a granulated nucleus and were positive for RNA polymerase II; spermatocytes I (SPC-I) showed positive expression for both *sycp3* and RNA polymerase II; round spermatids (SPD-r) and spermatozoa (SP) shared similar morphology under DAPI staining but differed in RNA polymerase II staining, with round spermatids being positive and spermatozoa negative. Between 50 and 100 cells were counted for each flow-sorted population, and only populations with an enrichment above 70% were considered for WGBS experiments.

### WGBS sample collection and library construction

Sorted cell populations were dissolved in homogenization buffer (20 mM Tris pH 8.0, 100 mM NaCl, 15 mM EDTA, 1% SDS, 0.5 mg/ml Proteinase K) at 55°C. Two Phenol/Chlorophorm/Isoamylalcohol (25:24:1, PCI) extractions were performed. DNA was precipitated using 1/5 volume of 4 M NH4Ac and 2.5 volumes of ice-cold absolute ethanol, and 1uL of linear acrylamide. Samples were incubated overnight at -20°C. The DNA was pelleted and resuspended in nuclease free water. Bisulfite converted libraries were generated using the Pico Methyl-Seq Library Prep Kit (Zymo Research, Cat. D5456) following manufacturer’s instructions. Libraries were sequenced on the Illumina NovaSeq X Plus Series platform (150 PE).

### WGBS analysis

Files were trimmed using Trimmomatic (v0.39) (HEADCROP:5 ILLUMINACLIP:TruSeq3-PE-2.fa:2:30:10 LEADING:3 TRAILING:3 SLIDINGWINDOW:4:15 MINLEN:50) ^100^ and mapped using Bismark (v0.22.3) (-- non_directional --local -X 2000) ^101^. Methylation was called using MethylDackel (v0.6.1) (--mergeContext -- minOppositeDepth 10 --maxVariantFrac 0.5). DMRs were detected using DSS (v2.54.0) (delta=0.2, p.threshold=0.05, minlen=100, minCG=10, dis.merge=100) ^102^.

## Acknowledgements

The Australian Research Council (ARC) Discovery Project (DP190103852); Ramón y Cajal fellowship (RYC2020-028685-I); and the Spanish Ministry of Science and Innovation project (PID2021-128358NA-I00), as well as funding from CEX2020-00108-M Unidad de Excelencia María de Maeztu to O.B. supported this work. J.J.T. was supported by the Spanish Ministry of Science and Innovation (PID2022-141288NB-I00). A.R-H. is funded by the Spanish Ministry of Science and Innovation (PID2020-112557GB-I00 funded by AEI/10.13039/501100011033 to A.R-H.), the Agència de Gestió d’Ajuts Universitaris i de Recerca, AGAUR (2021SGR00122 to A.R-H.) and the Catalan Institution for Research and Advanced Studies (ICREA). G.P. is supported by FPI predoctoral fellowships from the Ministry of Economy and Competitiveness (PRE-C-2021-0083). M.A-C. was supported by the Spanish Ministry of Science and Innovation (PID2020-113647GA-I00). The authors thank the Garvan Genomics Core for scATAC-seq and scRNA-seq library preparation, Nerea Roher for technical assistance, and members of O.B. and M.A-C. labs for critical reading of the manuscript.

## Contributions

O.B. and J.J.T. conceived the study. A.M.B.R. performed scRNA-seq and scATAC-seq data analyses with the input of J.J.T. and O.B. Zebrafish testis collection and processing (prior to library preparation) was performed by F-S.G. G.P. and A.R-H. performed zebrafish germ-cell sorting by FACS and immunofluorescence experiments. E.S. and M.A-C. generated testis cryosections and performed fluorescent *in situ* hybridization (FISH) experiments. O.B. and T.B. generated and analysed WGBS DNA methylome data. O.B. wrote the manuscript with the help of A.M.B.R. and J.J.T. All authors contributed to, read, and approved the final manuscript.

## Competing interests

The authors declare no competing interests.

## Notes

### Competing Interest Statement

The authors have declared no competing interest.

