## Supplementary Figures S1-S5 for "A single-cell multiomics roadmap of zebrafish spermatogenesis reveals regulatory principles of male germline formation"

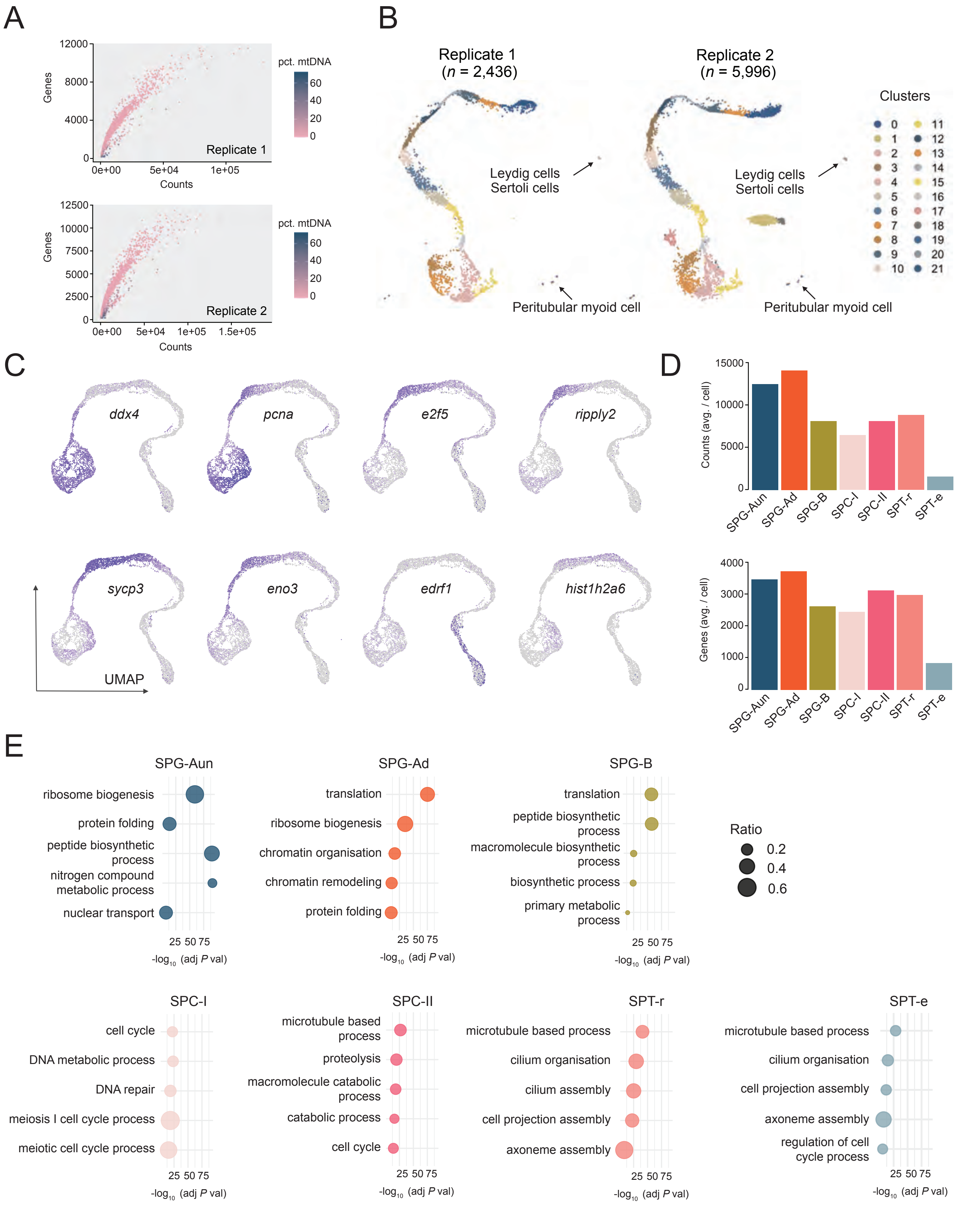


**Figure S1. scRNA-seq quality control, marker genes and GO enrichments. (A)** Quality control metrics for scRNA-seq replicates showing the relationship between counts, detected gene numbers, and mitochondrial DNA percentage. **(B)** UMAP (uniform manifold approximation and projection) plots of the zebrafish testis tissue denoted by Seurat clusters before filtering. **(C)** Distribution of average counts, and average gene numbers across the spermatogenic lineage. **(D)** Expression UMAP for canonical marker genes for SPG (*ddx4*), SPG-Aun (*eno3*, *e2f5* and *ripply2*), SPG-Ad(*hist1h2a6*), SPT-e (*edrf1*), SPC (*sycp3*) and proliferation (*pcna*). **(E)** Dot plot of the top 5 GO biological processes enriched using marker genes for each cell type. The x-axis represents adjusted *P* values, and dot size reflects the ratio of intersected to total terms.

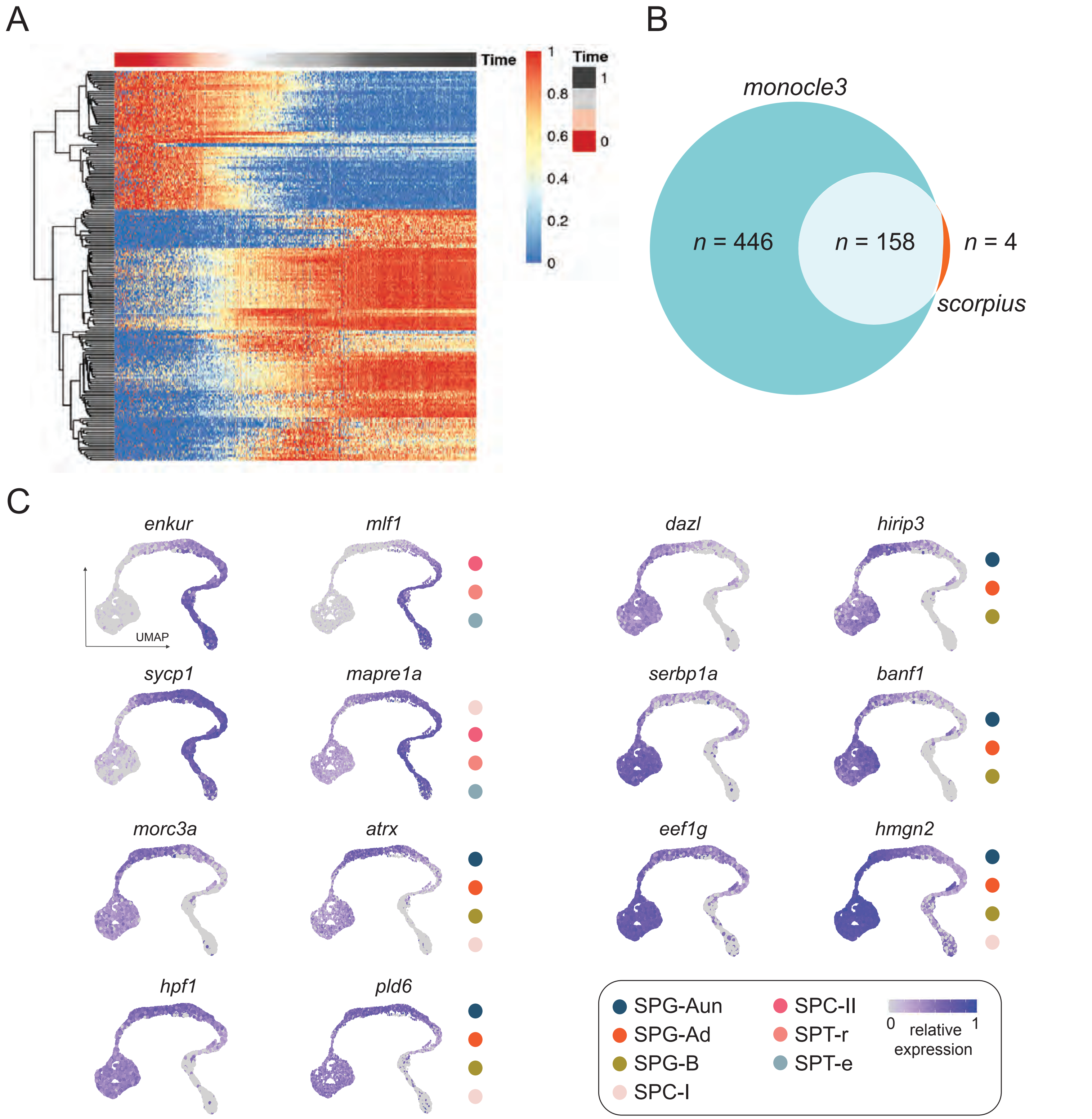


**Figure S2. Trajectory analysis and expression profiles of spermatogenesis driver genes. (A)** Heatmap of drive genes across the inferred trajectory of spermatogenesis. Samples ranked by pseudotime trajectory inferred by SCORPIUS. Rows represent genes grouped by hierarchical clustering with distinct expression patterns. **(B)** Comparison of gene drivers for spermatogenesis calculated using Monocle3 and SCORPIUS software. **(C)** UMAP examples for gene drivers obtained from the trajectory analysis identified by both Monocle3 and SCORPIUS.

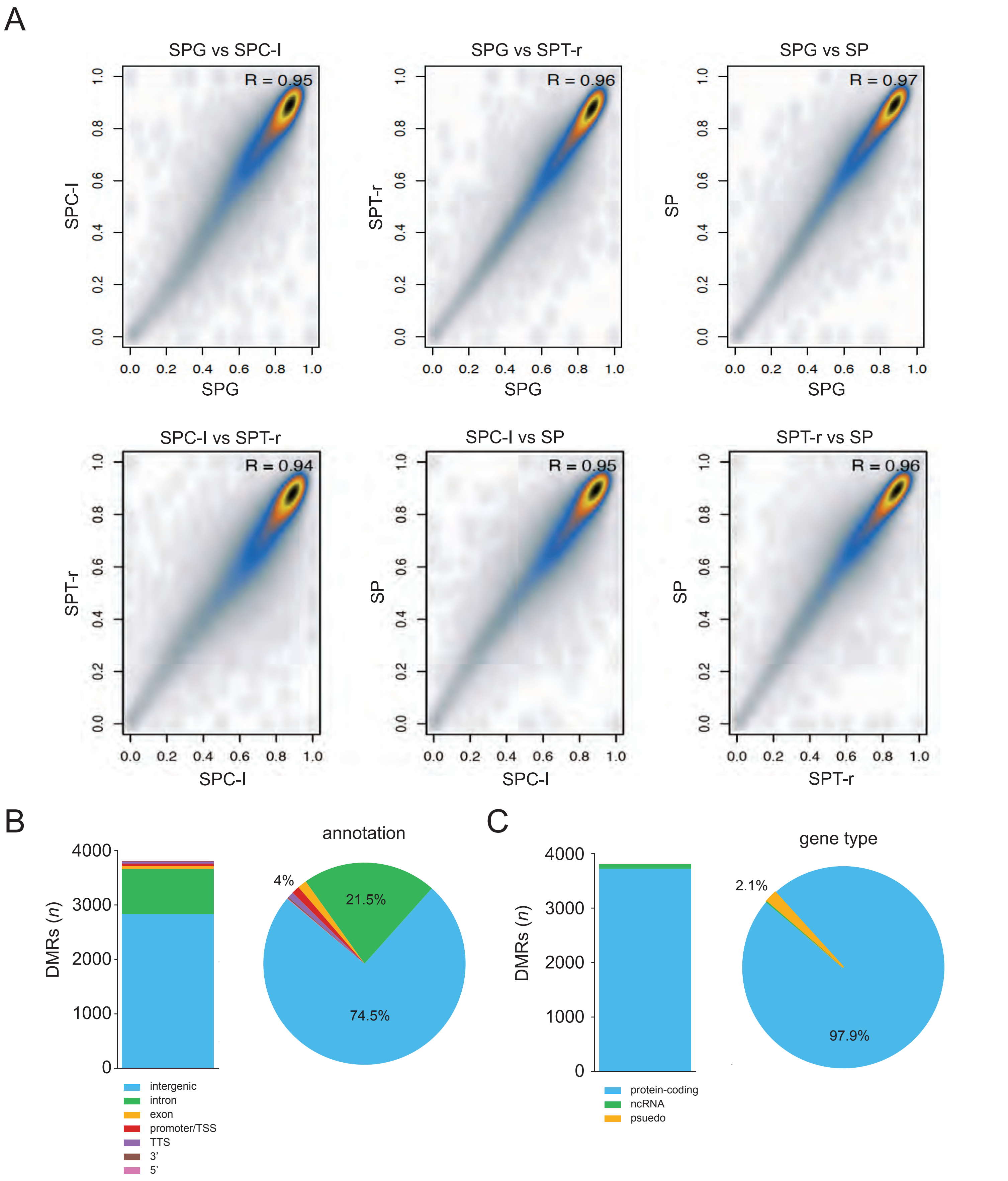


**Figure S3. DNA methylomes and genomic localisation of Differentially Methylated Regions (DMRs). (A)** Correlation of CpG methylation levels in non-overlapping 10 Kb genomic bins in isolated germ cell populations (SPG: spermatogonia, SPC-I: spermatocytes I, SPT-r: round spermatids, and mature sperm: SP). *R* value representes Pearson Correlation coefficient. **(B)** Genomic localisation represented as asbolute values (left panel) and percentages (pie chart, right panel) of DMRs (*n* = 3,879) identified between four germ cell populations. **(C)** Gene type enrichment of the identified DMRs.

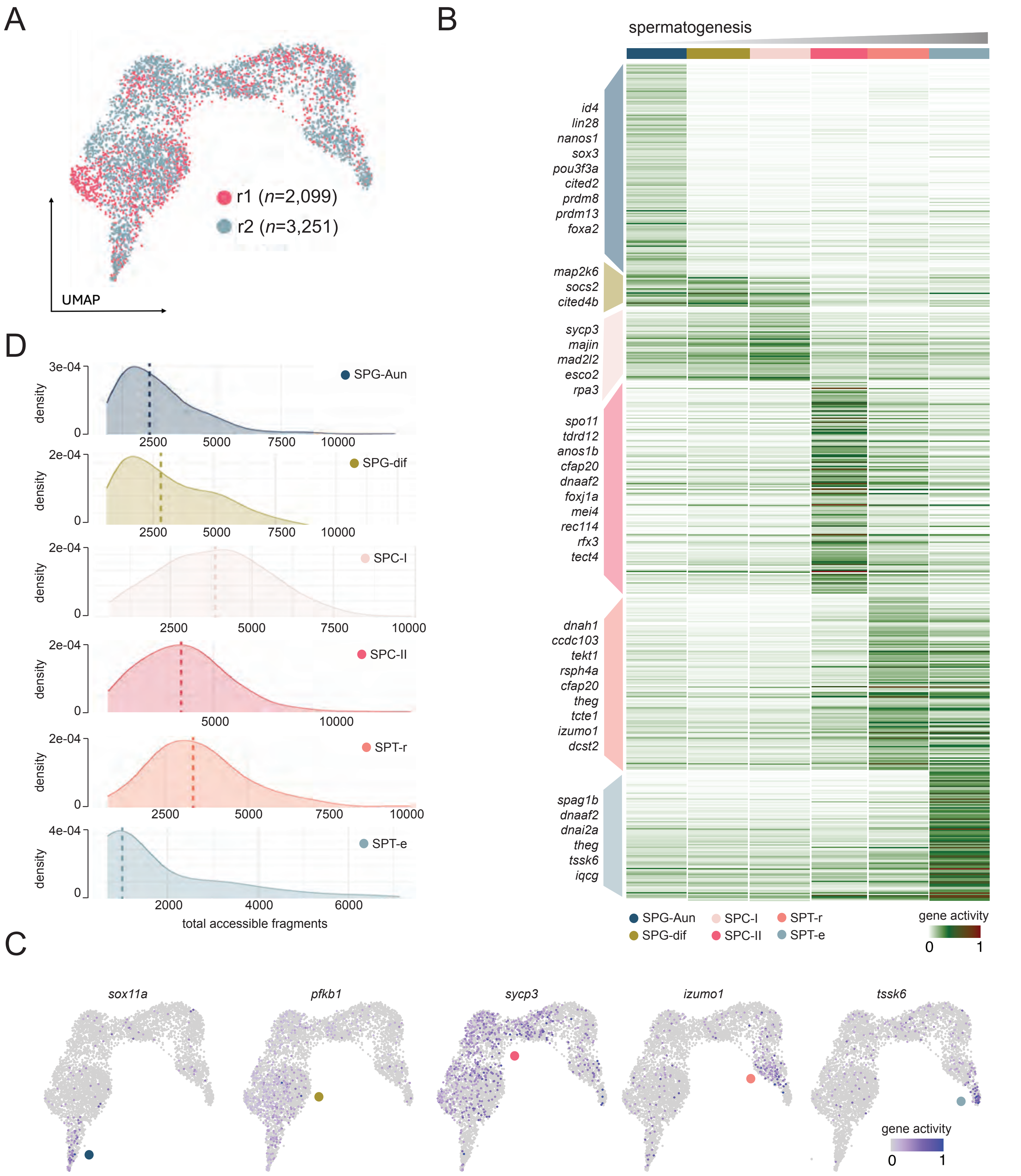


**Figure S4. Single cell chromatin accessibility (scATAC-seq) dynamics during zebrafish spermatogenesis. (A)** UMAP (uniform manifold approximation and projection) of biological replicates of scATAC-seq data. Cells are colored by replicate label. **(B)** Sorted heatmap of averaged gene expression profiles grouped by cell type (SPG-Aun, SPG-dif, SPC-I, SPC-II, SPT-r, SPT-e). Top marker genes for each cluster are indicated on the left hand side of the heatmap. **(C)** UMAP plots showing gene activity of marker gene examples associated with spermatogenesis. *sox11* - undifferentiated spermatogonia; *pfkb1* - differentiated spermatogonia; *sycp3* - spermatocytes; *izumo1* - round spermatids; *tssk6* - elongated spermatids. Gene activity is defined as the normalised sum of fragments spanning the gene body and extending 2 kb upstream of the transcription start site. **(D)** Distribution of the number of accessible genome fragments in each cell type (SPG-Aun, SPG-dif, SPC-I, SPC-II, SPT-r, SPT-e). The median is indicated by the dashed line.

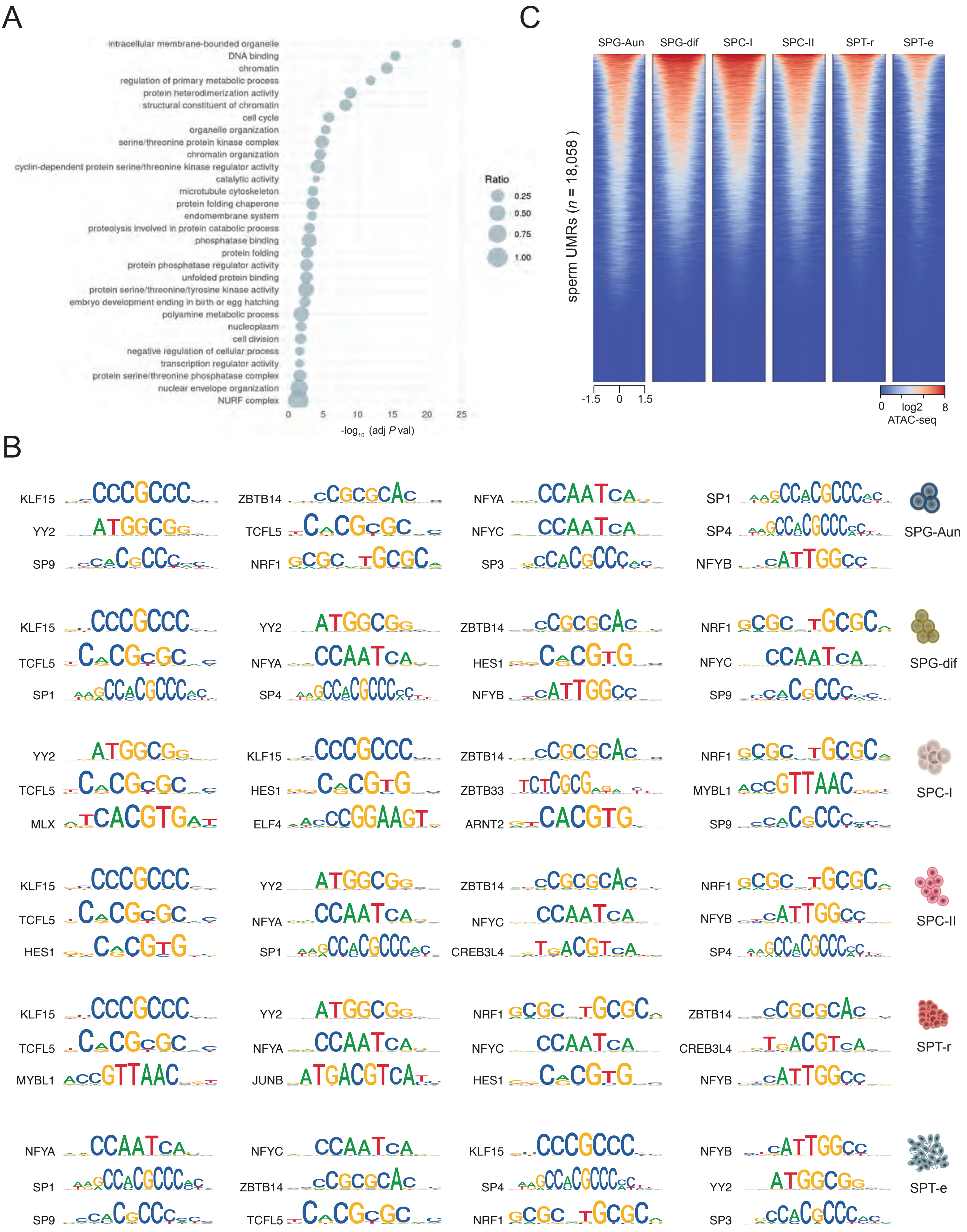


**Figure S5. Chromatin state of elongated spermatids (SPT-e). (A)** Dot plot of top biological processes associated with genes linked to open chromatin peaks in the SPT-e population. The x-axis represents adjusted *P* values, and dot size reflects the ratio of intersected to total terms. **(B)** Top 12 enriched motifs found in open chromatin peaks in each cell population **(C)** ATAC-seq signal (log2) from all spermatogenesis cell types, plotted over unmethylated regions (UMRs) identified in mature sperm.
